# Pseudotime trajectory analysis reveals divergent rod photoreceptor states during dark adaptation

**DOI:** 10.64898/2026.03.06.710240

**Authors:** Ryutaro Ishii

## Abstract

Rod photoreceptors face a high ATP demand in darkness, yet the molecular programs supporting dark adaptation remain difficult to dissect *in vivo*. Here, I re-analyzed publicly available murine retinal single-cell RNA-seq data and reconstructed rod-state dynamics across the light-to-dark transition using pseudotime trajectory inference. I found that dark-adapting rods diverge from a common state into two distinct lineages. Lineage 1 is characterized by elevated MYC-driven anabolic programs alongside increased reactive oxygen species (ROS) response and unfolded protein response (UPR)/ER stress signatures. In contrast, Lineage 2 exhibits an altered RNA-processing state with a markedly higher unspliced RNA fraction. Intronic motif analysis of Lineage 2 identified enriched binding sites for splicing-associated RNA-binding proteins linked to core spliceosome components and ATP-dependent helicases. Furthermore, Lineage 2 relatively preserves the STRADA/MO25β (CAB39L) module, which supports LKB1-AMPK energy sensing. Together, these findings support a model in which dark-adapting rods bifurcate into a MYC-driven anabolic state and an RNA-processing-altered state, a balance that may be biased by an LKB1–AMPK-associated mechanism. This integrative computational framework provides a predictive model for rod photoreceptor adaptation, guiding future biochemical and molecular experiments to determine the molecular basis and drivers of this divergence.

## Introduction

Rod photoreceptors (rods) consume substantially more ATP in darkness than in light and must rapidly adapt while maintaining cellular homeostasis [1]. This marked metabolic load can engage multiple stress pathways (including hypoxia-inducible factor (HIF) signaling and reactive oxygen species (ROS) production). A recent study suggested that extracellular adenosine may also influence ATP dynamics under dark conditions [2]. Because these pathways may operate simultaneously, identifying the key regulators coordinating this adaptation remains challenging. Dissecting these pathways *in vivo* often requires transgenic mouse models, which are time-consuming and not well suited to broad, unbiased screening across multiple candidate mechanisms. Data-driven approaches are therefore useful for prioritizing regulatory pathways for downstream validation.

Here, I re-analyzed publicly available murine retinal single-cell RNA sequencing (scRNA-seq) data [3] and reconstructed rod state dynamics across the light-to-dark transition using pseudotime trajectory analysis [4]. I found that dark-adapting rods diverged into two inferred lineages: one associated with MYC-linked anabolic programs and elevated ROS and UPR/ER-stress signatures, and the other characterized by a higher unspliced RNA fraction consistent with altered RNA-processing. Although the basis for this divergence remains unclear, these lineage-specific signatures suggest a link between energy sensing and post-transcriptional regulation during dark adaptation. These findings propose candidate regulatory mechanisms for future experimental validation to determine the basis and drivers of this divergence, as well as whether these states are reversible or stable.

## Materials and methods

### Data source and preprocessing

Publicly available retinal scRNA-seq data from adult wild-type mice (C57BL/6J, over 2 months old), obtained under normal light and dark-adapted conditions, were downloaded from the Genome Sequence Archive (GSA; accession CRA006518) [3]. Raw FASTQ files were reprocessed with Cell Ranger count (10x Genomics; version 10.0.0) using the mm10 reference genome. Ambient RNA contamination was reduced using SoupX [5], cells were filtered using standard quality-control metrics in Seurat (version 5.4.0) [6], and doublets were removed using DoubletFinder [7]. A whitelist of QC-passing singlets was then used to generate spliced and unspliced count matrices with velocyto (version 0.17.17) [8].

### Dimensionality reduction and clustering

After normalization and scaling, principal component analysis (PCA) was performed, and the first 15 principal components were used for UMAP visualization and Seurat graph-based clustering (FindNeighbors/FindClusters, resolution = 0.3). For rod-focused analysis, rod photoreceptors were computationally subset and re-clustered using the same parameters.

### Trajectory inference and pseudotime

Pseudotime trajectories were inferred with Slingshot (version 2.18.0) [4] using UMAP embeddings and graph-based cluster labels. Cluster 5, enriched for Light-condition cells, was designated as the root state. Pseudotime values were computed for individual cells along each inferred lineage.

### Unspliced RNA fraction analysis

Spliced and unspliced count matrices were imported into Seurat, where the single-cell unspliced fraction was calculated as unspliced / (spliced + unspliced) from the corresponding assay layers.

### Intron sequence extraction and RNA motif enrichment

Genes with elevated unspliced counts in the Lineage 2 terminal state were identified with Seurat FindMarkers using Bonferroni-adjusted P values and an effect-size threshold (avg_log2FC > 0.25). Intronic regions were defined from the mm10 annotation as the union of annotated introns for each gene, and the corresponding sequences were extracted for *de novo* motif discovery with STREME (version 5.5.9) [9]. Enriched motifs were compared against the CISBP-RNA database using Tomtom [10] to nominate candidate RNA-binding proteins.

### Gene set scoring and functional enrichment

Single-cell gene set scores were computed with UCell (version 2.10.1) [11]. Hallmark gene sets were obtained from MSigDB via msigdbr [12], and pathway activity was additionally inferred with PROGENy [13] where indicated. Gene Ontology (GO) enrichment analysis was performed with clusterProfiler [14] using Benjamini–Hochberg correction.

### Lineage-wise statistical comparisons

Terminal states were compared primarily as Cluster 3 (Lineage 1) versus Cluster 2 (Lineage 2). Two-sided Wilcoxon rank-sum tests were used for single-gene and module-score comparisons unless otherwise specified.

### Upstream regulator and PPI analysis

To identify candidate regulators of lineage-specific terminal states, transcription factor enrichment analysis was performed on the terminal gene signatures using the enrichR package [15] with the ChEA, ENCODE, and TRRUST [16] databases. To prioritize upstream regulators of the *Strada/Cab39l* module, ChIP-Atlas (mm10) [17] was queried for factors with promoter-proximal binding peaks at both loci. Protein–protein interaction (PPI) networks were evaluated with STRINGdb (version 12.0; Mus musculus) [18], and GO Biological Process enrichment of the resulting candidate set was assessed with clusterProfiler. To evaluate interactions with the core spliceosomal machinery, I defined a set of essential ATP-dependent RNA helicases and scaffolding proteins that drive the spliceosome cycle [19].

### Statistical analysis

All statistical analyses were performed in R (versions 4.3.0 and 4.5.2). For two-group comparisons, effect sizes were summarized as the area under the ROC curve (AUC) derived from the Mann–Whitney U statistic; AUC > 0.5 indicates higher values in Lineage 1, whereas AUC < 0.5 indicates higher values in Lineage 2. For GO enrichment, adjusted P < 0.05 was considered significant after Benjamini–Hochberg correction.

## Results

### Pseudotime trajectory analysis of the light-to-dark transition reveals two distinct rod transcriptional lineages

To characterize transcriptional changes in rods during dark adaptation, I re-analyzed murine retinal scRNA-seq data obtained under normal light (Light) and dark-adapted (Dark) conditions. UMAP visualization and graph-based clustering of the integrated dataset revealed transcriptomic heterogeneity within the rod population (Fig. 1A).

**Fig. 1.**
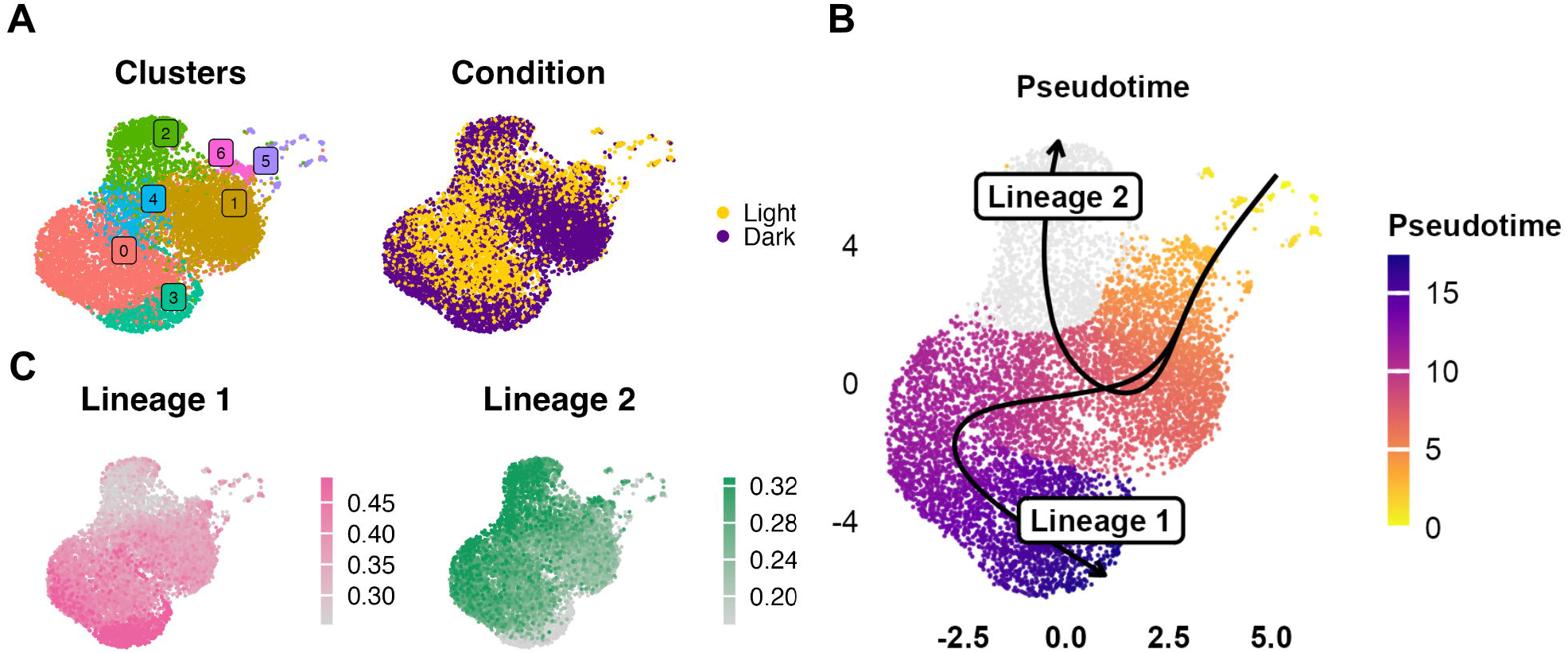
Pseudotime trajectory analysis reveals two lineages of rod photoreceptor states during dark adaptation. (A) UMAP visualization of re-clustered rod photoreceptor subpopulations (left) and the distribution of cells by experimental condition (Light vs. Dark) (right). (B) Slingshot pseudotime trajectory inference demonstrating the Lineage 1 path (Lineage 1: Clusters 5→6→1→4→0→3) and the Lineage 2 path (Lineage 2: Clusters 5→6→1→4→2). (C) UMAP projections showing UCell scores for the Lineage 1 (left) and Lineage 2 (right) gene signatures.

I next inferred a pseudotime trajectory using Slingshot, rooted at the Light-enriched starting state (Cluster 5), which identified a bifurcation from a shared starting state into two distinct lineages: Lineage 1 (Clusters 5→6→1→4→0→3) and Lineage 2 (Clusters 5→6→1→4→2) (Fig. 1B). To interpret the biological differences between the two terminal states (Cluster 3 for Lineage 1; Cluster 2 for Lineage 2), I derived lineage-specific terminal gene sets and computed UCell module scores, which showed largely non-overlapping patterns across the rod population (Fig. 1C). Gene Ontology enrichment further indicated distinct biological processes associated with each terminal state: Lineage 1 was enriched for cytoplasmic translation and oxidative phosphorylation, whereas Lineage 2 was enriched for RNA splicing-related pathways, cilium organization, and visual perception (Fig. S1). Collectively, these analyses suggest that rods undergoing light-to-dark adaptation diverge into two distinct transcriptional lineages with separable gene programs.

### Lineage 1 exhibits MYC-linked anabolic and stress-response signatures

To identify candidate upstream regulators, I integrated Enrichr-based transcription factor enrichment analysis of the lineage-specific terminal gene signatures with MSigDB Hallmark gene set scoring. MYC emerged as the top-ranked candidate transcription factor linked to the Lineage 1 terminal state (Fig. 2A). Hallmark scoring confirmed higher MYC target gene scores in Lineage 1 (AUC = 0.95) (Fig. 2B; Fig. S2A).

**Fig. 2.**
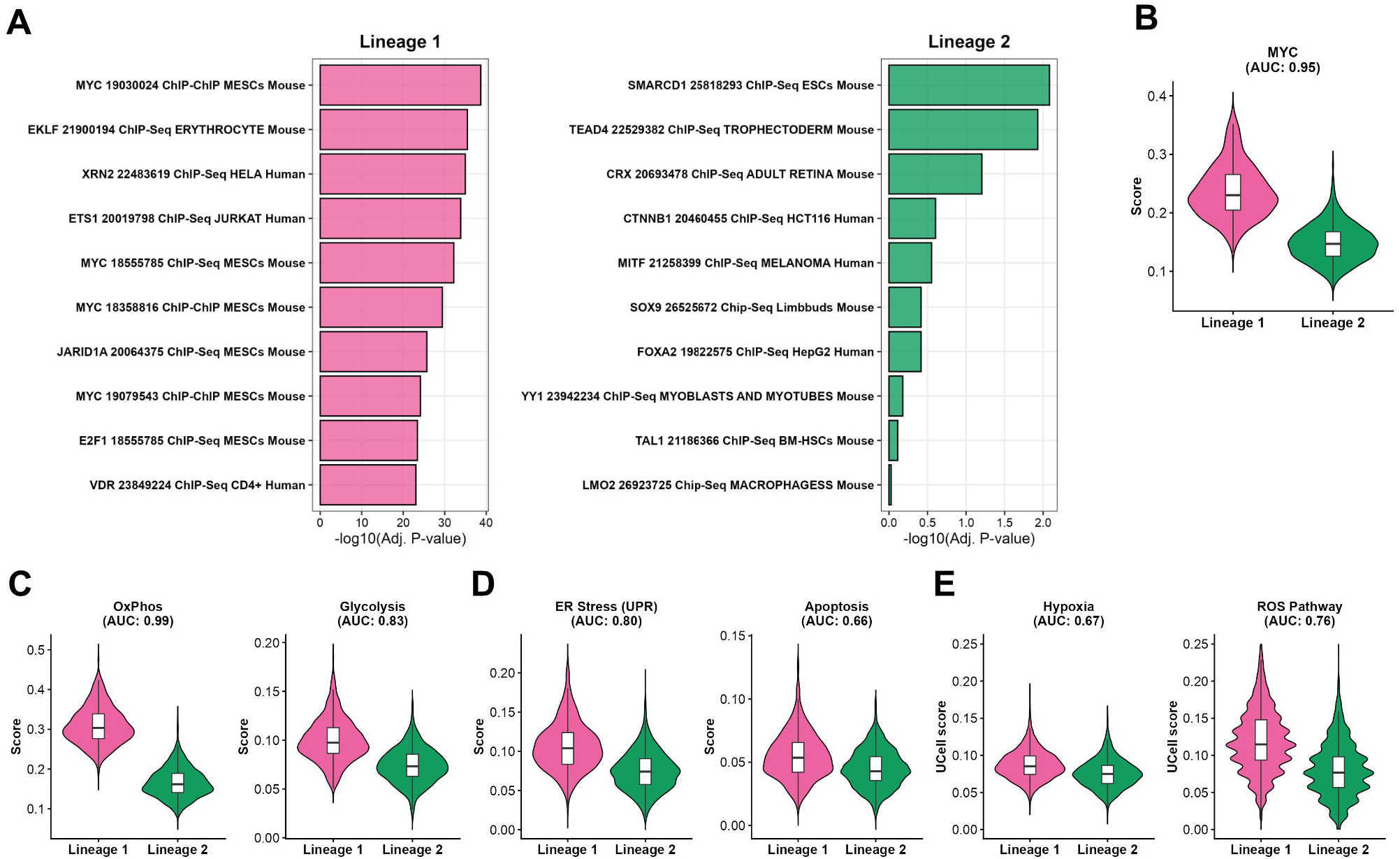
Lineage 1 exhibits enriched MYC-linked anabolic programs and stress-response signatures. (A) Data-driven enrichment analysis of upstream transcription factors using the ChEA and TRRUST databases. (B–E) Violin plots comparing MSigDB Hallmark gene set scores between Lineage 1 (magenta, left) and Lineage 2 (green, right) for MYC targets (B), oxidative phosphorylation and glycolysis (C), unfolded protein response (UPR) and apoptosis (D), and hypoxia and reactive oxygen species (ROS) pathways (E). Effect sizes are indicated by the area under the ROC curve (AUC; see Methods for details).

Because MYC is a master regulator of energy-consuming anabolic metabolism, I hypothesized that this lineage faces high energetic demands. Consistent with this MYC-associated profile, metabolic signatures for oxidative phosphorylation (AUC = 0.99) and glycolysis (AUC = 0.83) were prominently higher in Lineage 1 relative to Lineage 2 (Fig. 2C; Fig. S2B). Given that such elevated translation-associated programs can increase protein-folding demand, I next examined proteostasis-related signatures. The unfolded protein response (UPR)/ER stress signature was markedly higher in Lineage 1 (AUC = 0.80), whereas the apoptosis signature showed only a modest shift (AUC = 0.66) (Fig. 2D; Fig. S2C), suggesting increased proteostasis stress rather than immediate cell death.

I next examined stress-response signatures and inferred upstream signaling landscapes potentially sustaining this high-output state. The ROS response signature was higher in Lineage 1 (AUC = 0.76) (Fig. 2E; Fig. S2D). In PROGENy pathway inference, inferred EGFR pathway activity—a known upstream driver of anabolic networks—was the most increased in Lineage 1 among the evaluated pathways (AUC = 0.78) (Fig. S2E). In contrast, while the Hallmark hypoxia signature showed a modest increase in Lineage 1 (AUC = 0.67) (Fig. 2E; Fig. S2D), PROGENy-inferred hypoxia pathway activity was near neutral between lineages (AUC = 0.47) (Fig. S2E). Conversely, inferred TGF-β pathway activity was higher in Lineage 2 (AUC = 0.20; favoring Lineage 2) (Fig. S2E). These data indicate that Lineage 1 exhibits higher MYC-linked anabolic, ROS, and UPR/ER stress signatures than Lineage 2.

### Lineage 2 exhibits altered RNA-splicing signatures and accumulation of unspliced transcripts

Gene Ontology analysis of Lineage 2–enriched genes highlighted canonical rod functions. Notably, ‘RNA splicing’–related terms also emerged among the top-ranked categories (Fig. S1), prompting me to further examine post-transcriptional regulation. I therefore integrated spliced/unspliced mRNA profiling with intronic motif analysis and functional annotation. Cells in the Lineage 2 terminal state (Cluster 2) displayed a markedly higher unspliced fraction than those in the Lineage 1 terminal state (Cluster 3) (Fig. 3A, AUC = 0.07). Genes exhibiting elevated unspliced signals in Lineage 2 tended to be longer than those stalled in Lineage 1 (Fig. 3B, AUC = 0.33), consistent with altered processing kinetics that disproportionately affects long transcripts. Moreover, pathway annotation of these aberrantly retained transcripts revealed widespread intron retention specifically targeting genes essential for visual perception and photoreceptor development (e.g., *Sag, Prph2, Pde6b*) (Fig. 3C). To explore cis-regulatory features associated with the elevated unspliced signal, I performed *de novo* motif enrichment on intronic sequences using STREME. This identified five enriched motifs (Fig. 3D). Motif similarity searches with Tomtom suggested that these enriched motifs broadly resemble the binding preferences of known splicing-associated RBPs, including SR-family factors (e.g., SRSF7), RBM-family proteins (e.g., RBM45 and RBM42), and polypyrimidine/U-rich binding proteins (e.g., PTBP1, BRUNOL4). These candidate RBPs encompass diverse functional roles in splicing regulation, including enhancers, modulators, and silencers. One enriched motif (Motif 4; UGGCUCC) did not match a known RBP motif in the queried database.

**Fig. 3.**
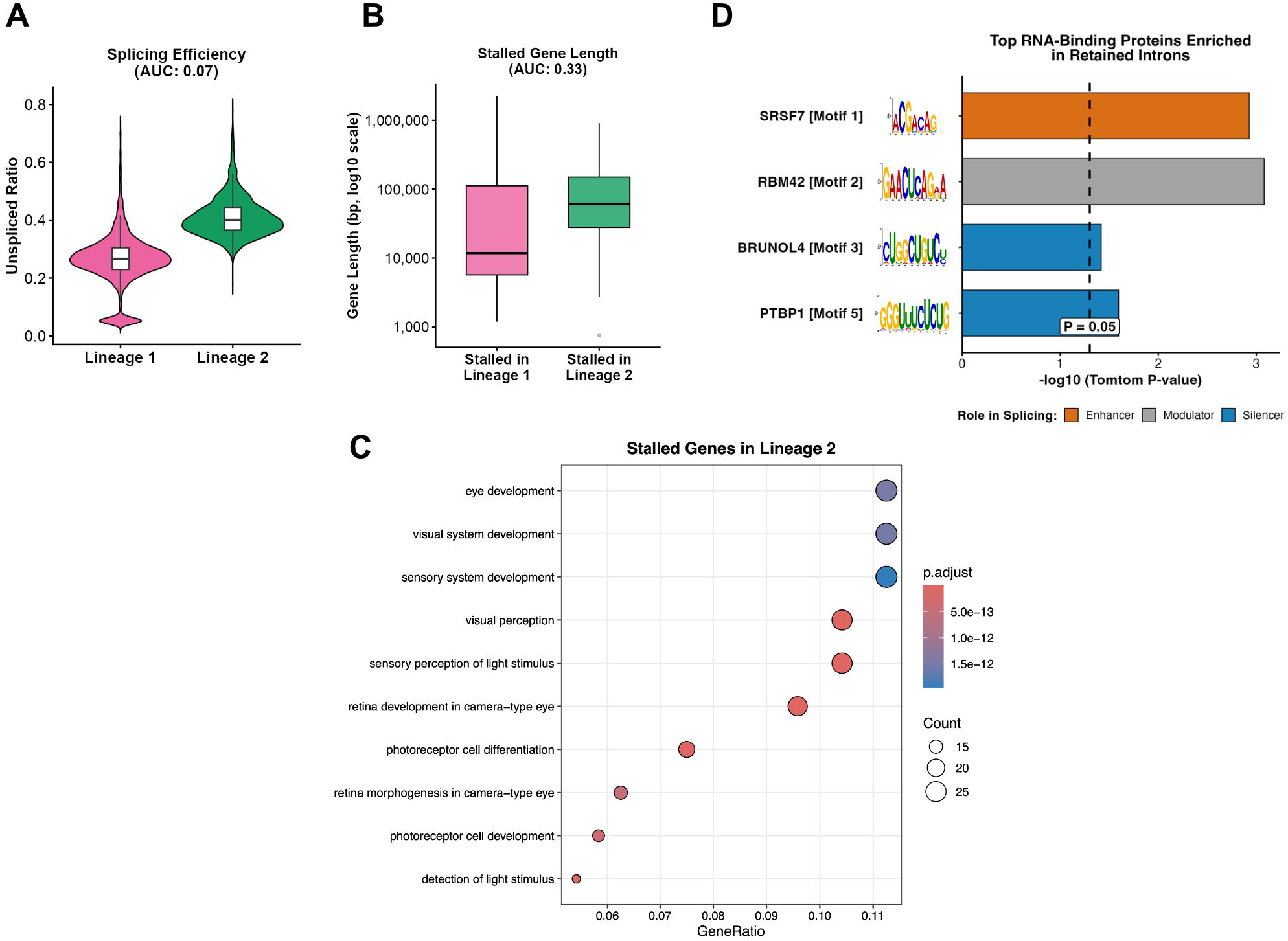
Lineage 2 shows increased unspliced RNA accumulation in long, vision-related transcripts, consistent with altered RNA-processing kinetics. (A) Comparison of the unspliced fraction between lineages (colors and statistical metrics as in Fig. 2), showing the accumulation of unspliced RNA signals in Lineage 2. (B) Genomic span of the genes exhibiting elevated unspliced signals in Lineage 2. (C) Gene Ontology (GO) enrichment analysis of genes with elevated unspliced signals in Lineage 2, highlighting enrichment of terms related to visual perception and photoreceptor development. (D) *De novo* motif enrichment analysis of intronic sequences from genes with elevated unspliced signals using STREME, and motif similarity matching to known RNA-binding protein (RBP) motifs using Tomtom. The bar plot highlights selected representative splicing-associated RBPs matching the discovered motifs, categorized by their general roles in splicing regulation.

Because motif similarity alone does not establish mechanism, I next asked whether these candidate RBPs have known connections to core spliceosomal components. A STRING-based protein–protein interaction (PPI) analysis indicated that several candidates, including RBM42, are linked to core spliceosome proteins (e.g., PRPF8, EFTUD2) as well as ATP-dependent helicases (e.g., DDX23, SNRNP200) (Fig. S3). Together, these results associate the Lineage 2 trajectory with increased unspliced pre-mRNA accumulation and intronic sequence features suggestive of splicing-associated RBPs.

### Relative preservation of an LKB1–AMPK-associated module in Lineage 2

Given the extreme energetic demand placed on rods in darkness, I asked whether transcriptomic footprints of energy-sensing pathways differ between the two lineages. Consistent with the earlier findings of MYC-linked metabolic activation, the HALLMARK_MTORC1_SIGNALING score was higher in Lineage 1 (Fig. 4A; AUC = 0.88; Fig. S4A), consistent with sustained anabolic/translation-associated gene programs. In contrast, UCell scoring based on the KEGG AMPK signaling pathway was near neutral, with a slight tendency toward higher AMPK-associated gene-set scores in Lineage 2 (Fig. 4B; AUC = 0.48; favoring Lineage 2; Fig. S4B). Because KEGG/UCell scores reflect transcript abundance rather than direct kinase activity, I next examined the expression of key pathway components. Although the overall KEGG AMPK signature was near neutral, individual components showed lineage-associated differences. Within the KEGG AMPK pathway gene set, Lineage 1 exhibited higher expression of *Adipor1* and *Tsc2*, whereas *Strada* and *Cab39l* (MO25β)—components of the LKB1–AMPK activating complex—were relatively higher/maintained in Lineage 2 (Fig. 4C). Together, these lineage-associated differences highlighted the STRADA/MO25β-LKB1-AMPK module for further upstream-regulator analysis.

**Fig. 4.**
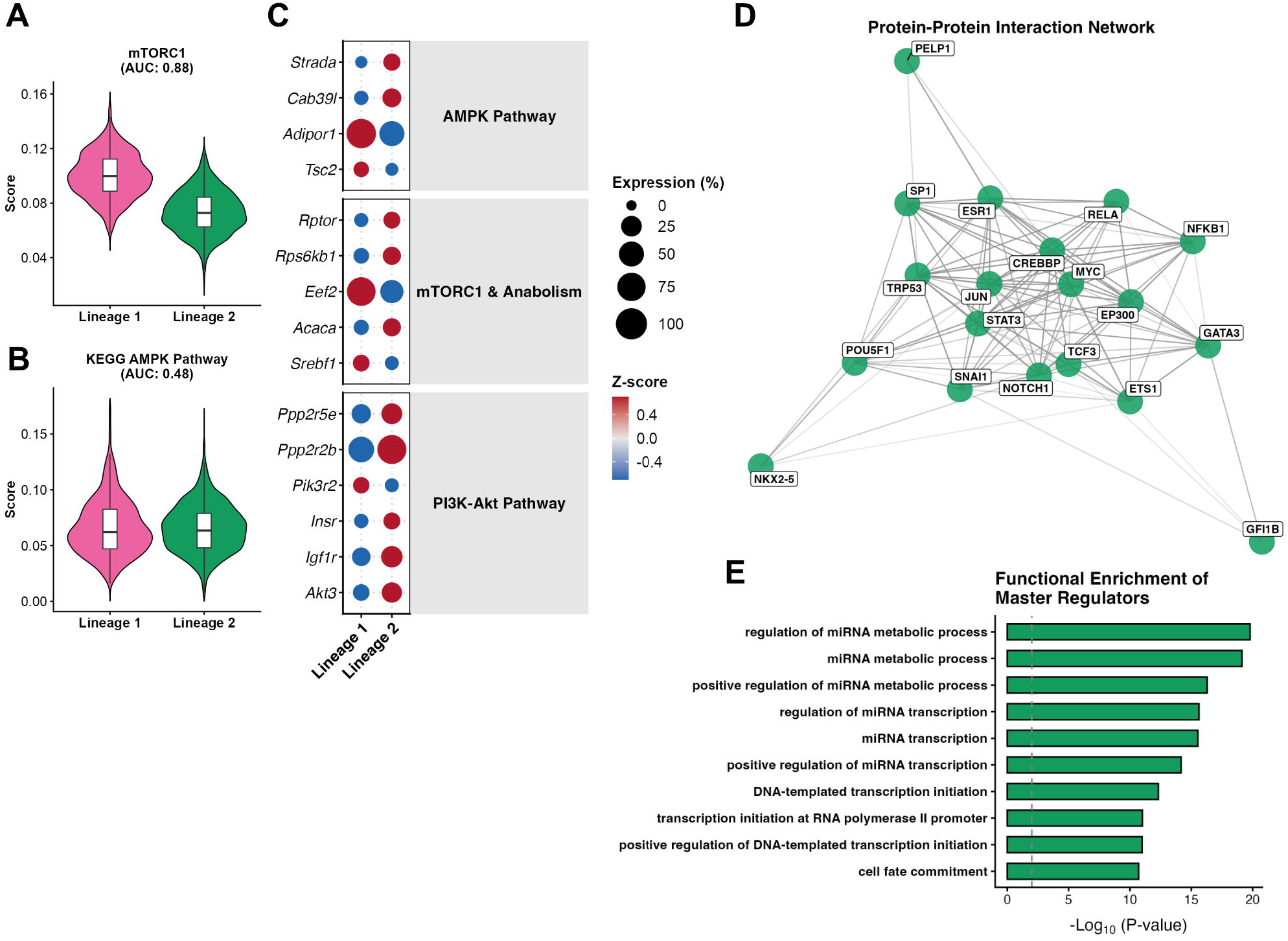
Differential mTORC1- and AMPK-associated transcriptional footprints across lineage divergence nominate candidate regulators of the STRADA/MO25β (CAB39L). (A, B) Distribution of UCell scores for the MSigDB Hallmark mTORC1 signaling gene set (A) and the KEGG AMPK signaling pathway gene set (B) in Lineage 1 (left, magenta) and Lineage 2 (right, green). (C) Dot plot of differentially expressed genes (DEGs) between Cluster 3 (Lineage 1) and Cluster 2 (Lineage 2), filtered to genes in the KEGG AMPK signaling pathway. (D) Interaction network of candidate upstream regulators nominated from public ChIP-seq compendia (ChIP-Atlas; transcription factors with promoter-proximal binding evidence near both *Strada* and *Cab39l*) integrated with TRRUST annotations. Edges indicate known or predicted functional associations from the STRING database. (E) Gene Ontology (GO) Biological Process enrichment analysis of the core master regulatory module identified in (D). The bar plot displays the top statistically enriched functional terms (calculated via clusterProfiler), highlighting the module’s profound involvement in the regulation of miRNA metabolic processes.

To generate hypotheses for upstream regulators linked to *Strada/Cab39l* expression, I queried public ChIP-seq compendia (ChIP-Atlas) to identify transcription factors reported to bind near the promoter regions of both genes, and integrated these candidates with TRRUST-based regulatory annotations. This prioritization yielded a focused set of candidate regulators. Protein–protein interaction analysis further suggested that many of these candidates are connected within a shared interaction network (Fig. 4D). Gene Ontology enrichment analysis of this candidate set revealed enrichment for terms related to miRNA metabolic processes (Fig. 4E).

## Discussion

In this study, I re-analyzed publicly available murine retinal scRNA-seq data and reconstructed rod-state dynamics during dark adaptation using pseudotime trajectory inference, revealing a divergence into two distinct lineages. Lineage 1 represents a MYC-linked high-output anabolic state accompanied by elevated ROS and UPR signatures. These stress signatures have been implicated in retinal degeneration [20,21], and the higher mTORC1-associated signature in Lineage 1 is consistent with prior evidence linking insulin/mTOR signaling to retinal degeneration [22]. This raises the possibility that Lineage 1 cells may be less able to attenuate energetic and translational demands under stress, which could contribute to the accumulation of oxidative and proteotoxic stress. By contrast, Lineage 2 represents an altered RNA-processing state in which unspliced RNAs accumulate and translation output could potentially be reduced via post-transcriptional mechanisms.

Consistent with this, intronic motif analysis revealed an enrichment of binding sites for splicing-associated RBPs, and network analysis linked these candidates to core spliceosome components. Together, my current data support a working model of Lineage 2 in which splicing regulation may contribute to metabolic adaptation of rods, potentially through interactions between intrinsic intronic features of vision-related genes and the activity/composition of the splicing machinery.

Given that several candidates connect to ATP-dependent helicases in the spliceosome, I next asked whether energy-sensing pathways also differ across the divergence. Lineage 1 showed higher mTORC1-associated signatures, whereas Lineage 2 relatively preserved the expression of *Strada* and *Cab39l* (encoding STRADA and MO25β), which form the LKB1-activating complex required for efficient AMPK activation [23]. Although my transcriptomic data do not directly measure kinase activity, these results raise the possibility that the capacity to engage the LKB1–AMPK checkpoint may differ between the two lineages. Normally, AMPK is activated by energy depletion or oxidative stress and suppresses ATP-consuming anabolic pathways, including mTORC1-driven protein synthesis [24]. In this framework, reduced expression of the STRADA/MO25β module in Lineage 1 could indicate a lower capacity to support LKB1-dependent AMPK activation, potentially helping to explain why mTORC1-linked signatures persist in Lineage 1 despite elevated stress signatures.

Furthermore, candidate upstream regulators of *Strada* and *Cab39l* were enriched for miRNA metabolic processes, nominating miRNA-mediated post-transcriptional regulation as an additional layer that could tune this energy-sensing module. Although the biological relevance of this mechanism requires experimental validation, MYC can interface with miRNA regulatory programs [25] and retained-intron transcripts have been proposed to potentially bind multiple miRNAs and inhibit their original functions [26]. Indeed, miRNAs are known to function in light/dark adaptation [27], and miRNA-dependent gene silencing via the RNA-induced silencing complex (RISC) is essential for retinal homeostasis and stress responses [28]. Moreover, my PROGENy analysis inferred higher TGF-β pathway activity in Lineage 2, consistent with a recent pseudotime analysis in a retinal light-damage model that reported regulation of TGFβ signaling and the RISC complex [29]. Although these contexts differ from dark adaptation, these reports support my miRNA hypothesis. Together, these data highlight a previously unrecognized emergence of bifurcating rod states during dark adaptation, which might be regulated by the interaction of RNA processing, miRNA/RISC regulation, and the LKB1–AMPK module. To move beyond current transcriptomic correlations, future studies should employ biochemical assays of AMPK–mTORC1 signaling, as well as direct measurements of spliceosomal assembly and intron retention alongside gene-expression profiling and miRNA/RISC regulation. These studies will be essential to determine why two distinct lineages emerge during dark adaptation, what the drivers are that determine these lineages, and whether they represent reversible states or stable endpoints.

## Author contributions

R.I.: Conceptualization, Methodology, Formal analysis, Investigation, Data curation, Visualization, Writing – original draft, Writing – review & editing.

## Acknowledgements

I thank Dr. Hiromi Yanagisawa for generously providing access to the computational resources essential for my data analysis. I would like to acknowledge the authors of the original study (GSA accession CRA006518) for making their invaluable scRNA-seq dataset publicly available. During the preparation of this work, the author used Generative AI assistance (Gemini, Google) in order to help draft and debug custom analysis scripts, as well as to assist with English language editing and stylistic improvements. After using this tool/service, the author reviewed and edited the content as needed and takes full responsibility for the content of the published article.

## Data Availability Statement

The scRNA-seq data analyzed in this study are publicly available from the Genome Sequence Archive (GSA accession CRA006518). Custom analysis scripts used in this study are available from the corresponding author upon reasonable request.

