## Supplementary figures for "Pseudotime trajectory analysis reveals divergent rod photoreceptor states during dark adaptation"

Fig. S1

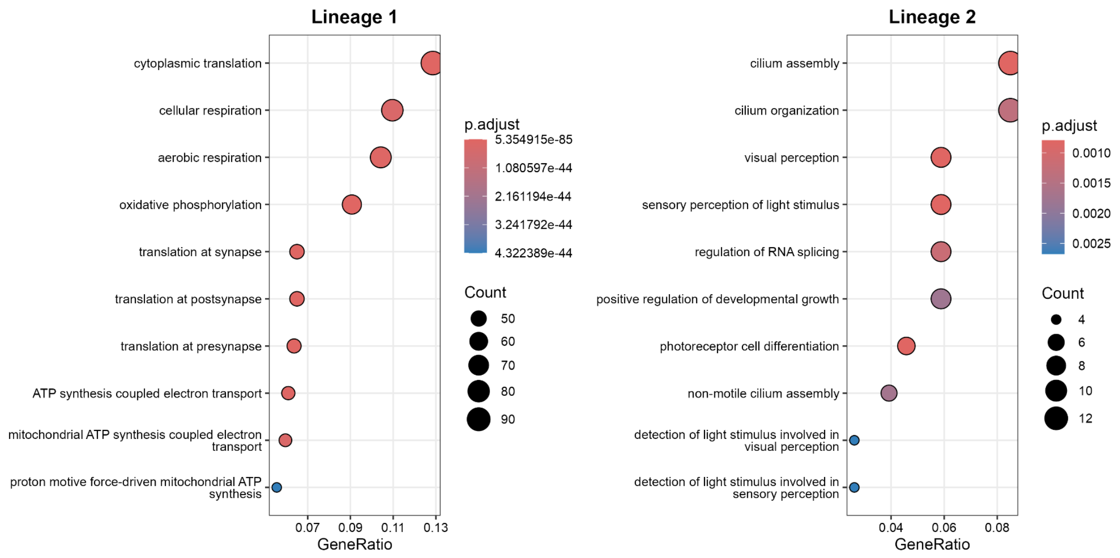


**Fig. S1. Gene Ontology enrichment analysis of rod lineages.**

Gene Ontology (GO) enrichment analysis of late-stage gene signatures specific to Lineage 1 (Cluster 3) and Lineage 2 (Cluster 2).

**Fig. S2**

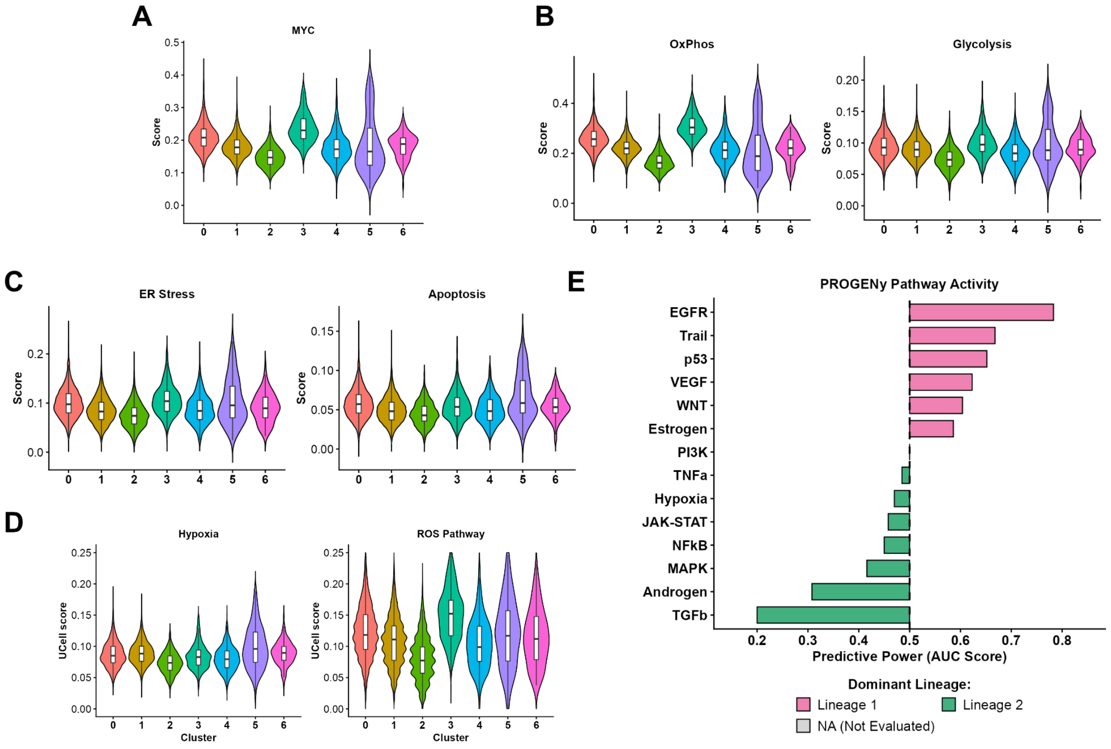


**Fig. S2. MSigDB Hallmark gene set scores and PROGENy pathway activity across clusters.**

(A–D) Violin plots showing MSigDB Hallmark gene set scores across all clusters for MYC targets (A), Oxidative Phosphorylation and Glycolysis (B), ER stress (Unfolded Protein Response) and Apoptosis (C), and Hypoxia and Reactive Oxygen Species (ROS) pathways (D). (E) Diverging bar plot showing PROGENy pathway activity comparing Cluster 3 (Lineage 1) and Cluster 2 (Lineage 2).

Fig. S3

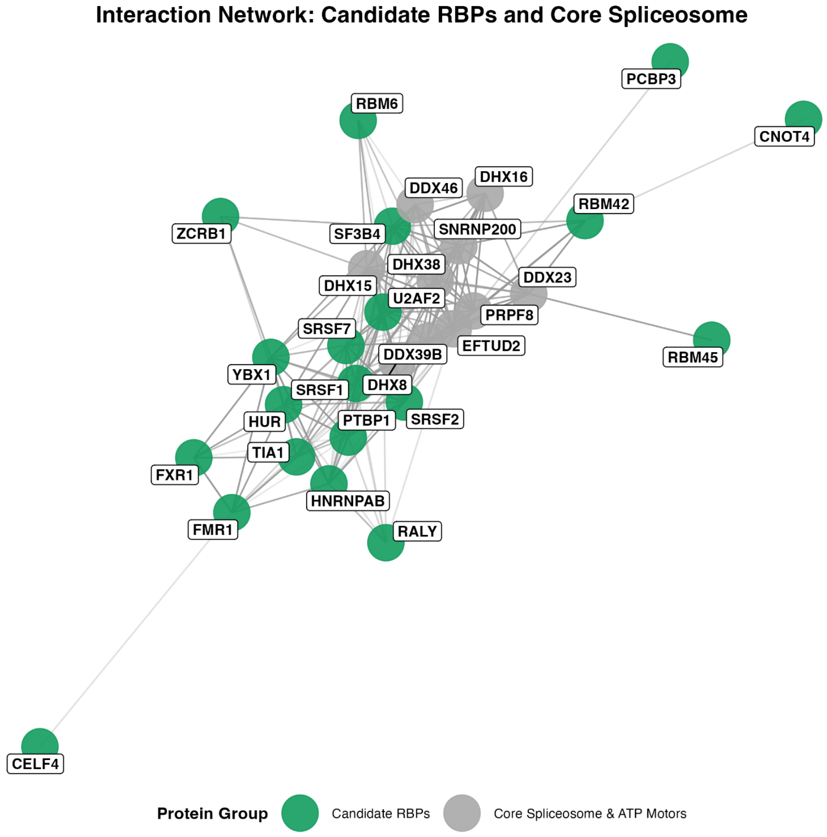


**Fig. S3. Protein–protein interaction network.**

STRING-based protein–protein interaction (PPI) network analysis. The network illustrates physical and functional associations between the identified candidate RNA-binding proteins (e.g., RBM42; green nodes) and core spliceosome machinery, including ATP-dependent RNA helicases. Core spliceosome components and ATP-dependent helicases (dark gray nodes) were selected based on the highly conserved spliceosomal key players (DDX46, DDX23, DDX39B, SNRNP200, DHX16, DHX38, DHX8, DHX15, PRPF8, and EFTUD2). Further details are provided in the Materials and methods section.

**Fig. S4**

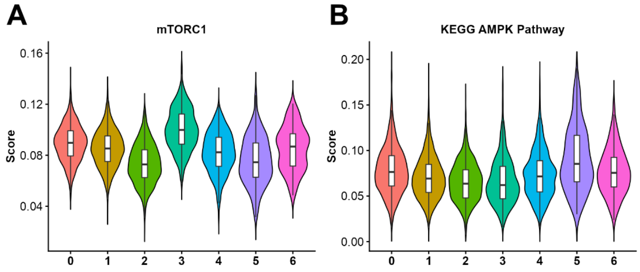


**Fig. S4. Extended expression profiles of energy-sensing pathways.**

(A, B) Violin plots detailing the distribution of UCell scores for the MSigDB Hallmark mTORC1 signaling gene set (A) and the KEGG AMPK signaling pathway (B) across all identified clusters.
